# A spin glass representation of hierarchical gene regulation and the partial resolution of frustration

**DOI:** 10.64898/2026.10.29.762975

**Authors:** Taylor E. Woodward, Michael P. Stumpf

## Abstract

Gene regulatory networks are directed and often contain loops. A loop can place competing demands on its genes, so that no expression state satisfies every regulatory interaction. In statistical physics such a conflict is called frustration. Existing spin-glass models of tissues assume symmetric regulation, and studies of frustration consider single cells. We develop a nonreciprocal multicellular Ising model that couples a directed regulatory network in each cell to random, symmetric signalling between neighbouring cells. We define intracellular and intercellular frustration densities, which count the unsatisfied interactions within a cell and with its neighbours. We study six exemplar nine-gene networks that share a downstream cascade and differ in an upstream three-gene motif. In isolated cells the motif sets the frustration. Negative feedback loops pass frustration around the loop, and their genes keep switching. The incoherent feedforward loop holds frustration at a single gene and freezes. When cells communicate, they form spatial clusters with shared expression profiles, and conflict between neighbours concentrates at cluster boundaries. Stronger coupling lowers intercellular frustration and raises intracellular frustration. Frustration is therefore resolved only in part: cells satisfy more of the couplings to their neighbours by satisfying fewer of their own regulatory interactions. Varying the coupling strength reveals a transition near the value at which the combined input of a cell’s eight neighbours matches a single regulatory interaction. Below it, cells follow their own networks and the tissue fragments. Our framework links the structure of gene regulatory networks to the stability of gene expression in cells and tissues.

## 1 Introduction

The differential regulation of genes is known to be one of the crucial factors determining cellular physiology [1, 2]. It is also more closely linked to organismic complexity than e.g. gene number [1, 3]. We have a solid understanding of the molecular machinery involved in gene regulation as well as mRNA and protein expression [4, 5].

In parallel with the development of exquisite experimental technologies [6–8], the theoretical investigation into gene expression regulation has also become increasingly rich and rewarding. Reward here can come in two forms: first, better quantitative models that allow us to drill down into the kinetics of e.g. transcriptional initiation, the kinetics of transcription into mRNA, and the translation into protein [9–12]. Second, better and more comprehensive insights into regulatory logic behind the transcriptional programs [13] that accommodate, and arguably drive, cellular processes, ranging from cell-fate decision making [14–16], to stress response [17–19].

Despite the vast improvements that we have seen in these quantitative models, there is still scope and need for better qualitative understanding of fundamental problems in cell biology. One example that we discuss here is how cells reconcile different, potentially competing demands. We tackle this from as flexible and general a framework as seems reasonable and we use a model from statistical physics, which applies to the simple case where genes are either “ON” or “OFF” [20–23]. This can be the case for naturally evolved systems, but similar processes have also been generated synthetically. Discrete, all-or-none gene expression has been shown using e.g. single-cell flow cytometry [24], time-lapse microscopy [25–27], single-molecule FISH [28], and hysteresis experiments [29–31]. These have shown that many natural circuits (lac operon [25, 29, 30], lambda phage [26, 32], *Bacillus subtilis* competence [28, 33], yeast GAL [24, 31], chromatin/epigenetic binary switching [27, 34, 35], and HOX gene expression [36]) and engineered circuits (the toggle switch [37], recombinase memory devices [38, 39], CRISPRoff [40]) occupy two stable states instead of a continuum.

We will investigate dynamics here from the perspective of conflicting regulatory signals. In statistical physics conflicting inputs into e.g. a classical spin system or Boolean variable, give rise to something we refer to as frustration [41–44]. Below we develop that notion and show how frustration arises naturally in the context of gene regulatory networks [45–48]. We show the extent to which frustration affects very simple exemplar systems of gene regulation; and we discuss the extent to which cell-cell interactions modulate intra-cellularly conflicting results [49, 50].

## 2 Model and Methods

This section sets out the model and the measures we use. We first review the Ising model, spin glasses, and frustration, and then adapt a multicellular spin-glass model to directed gene regulation. Directed interactions break the symmetry of the model, and the system is no longer described by an equilibrium distribution. We then define metrics for the frustration of a cell, introduce the example gene regulatory networks that we study, and summarize the simulations.

### 2.1 Ising models, spin glasses, and frustration

We describe the state of each gene by a binary variable. The Ising Model, as defined in statistical physics, describes variables, such as spins, with states *σ*_*k*_ ∈ {−1, +1}, and here they represent the OFF and ON states of a gene [51]. The model was introduced to describe interacting magnetic spins on a lattice [52] and has since been applied in fields ranging from machine learning to cellular differentiation [53]. For symmetric couplings the energy of a configuration is given by the Hamiltonian

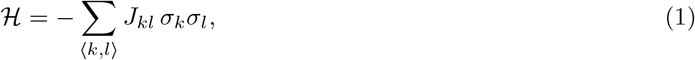

where *J*_*kl*_ = *J*_*lk*_ is the coupling between sites *k* and *l*, and the sum runs over interacting pairs, for example nearest neighbours on a lattice. Here *k* and *l* index sites; from Section 2.3 on, *i* and *j* index cells and *A* and *B* index genes. Configurations of low energy satisfy most couplings, and the minima of H are the stable states of the system. The model shows how local interactions give rise to global order, which makes it a general framework for emergent behaviour [54, 55].

Spin glasses extend the Ising model by assigning random signs to the couplings. Their energy landscapes have many local minima, which reflect competing and often contradictory interactions between components [56]. This suits biological systems, where the number of possible states is vast but the number of biologically relevant stable states is small [57, 58]. With the resolution of single-cell RNA sequencing, principles of statistical mechanics have been used to define the microstates and macrostates of cells in the context of cell fate [51, 57, 59, 60].

Competing interactions give rise to what is known as frustration. A system is frustrated when no configuration of its spins satisfies all of its couplings [44, 47, 55, 58]. The canonical example is a triangle of spins with antiferromagnetic couplings, which favour anti-alignment. Any configuration of the three spins leaves at least one pair aligned, so one coupling is always unsatisfied.

### 2.2 Directed gene regulation and nonreciprocal interactions

Gene regulatory networks (GRNs) are directed and often contain loops. A loop can place competing demands on its genes, whose expression states may then fluctuate. Such fluctuations may be noise, or a feature of cell development [47, 61]. Cell types are commonly identified with distinct gene expression profiles [51], and a GRN with competing interactions can produce more than one profile, and hence more than one cell type. Relating network structure to the stability of gene expression and of tissues therefore requires a model that keeps the direction of each interaction, *K*_*AB*_ ≠ *K*_*BA*_ [62] (Section 2.3).

Frustration is a key property of biological regulatory networks. Tripathi et al. model directed GRNs as asymmetric spin glasses on a graph, in which each gene follows the sign of its regulatory input [47, 48]. They find that minimally frustrated stable states correspond to cell types [47, 48, 63], while highly frustrated states are associated with ambiguous or disease-related phenotypes. These studies consider the GRN of a single cell in isolation, without communication between cells.

Incoherent feedforward loops (ICFFLs) are common motifs in GRNs. They play a role in the differentiation and proliferation of pluripotent stem cells [64, 65]. In an ICFFL the two paths from the input gene to the target gene have opposite signs, so the target receives conflicting signals. In a feedback loop, an odd number of repressive edges makes the loop negative, and in isolation a negative feedback loop oscillates and is frustrated [66]. Loops that share a node, known as conjugated loops, can reduce or remove the frustration of the network as a whole [45, 66]. Communication between cells can create such shared loops across cells. This motivates our question of how cell-cell communication changes the frustration that a loop motif creates within a cell.

Spin-spin interactions are usually taken to be symmetric. Gene regulatory interactions are not, and for directed couplings *J*_*kl*_ ≠ *J*_*lk*_. With asymmetric couplings no global energy function exists: single-spin-flip dynamics break detailed balance, and the steady state is no longer a Boltzmann distribution. Each spin lowers its own selfish energy, and when these goals conflict the system can cycle indefinitely [67]. Asymmetric couplings have a long history. Asymmetric spin glasses and neural networks were studied in the late 1980s [68, 69], and kinetic Ising models with asymmetric couplings are a standard tool for inferring connectivity from neural recordings [70]. Recent work has examined how nonreciprocal couplings change the collective behaviour of Ising models [67, 71]. These studies provide the basis for a nonreciprocal Ising model of directed gene regulation.

We therefore develop a nonreciprocal multicellular Ising model. It is a spin-glass model with an asymmetric intracellular interaction matrix, coupled to random, symmetric communication between neighbouring cells.

### 2.3 Nonreciprocal multicellular Ising model

Cells in a tissue interact through molecular signalling [47, 49]. Smart and Zilman built a two-level spin-glass model of tissue structure, in which the stochastic dynamics of spins model stochastic gene regulation and cell-cell communication [49]. The first level, which we call the **intracellular layer**, describes gene-gene interactions within a cell. The second, the **intercellular layer**, describes communication between neighbouring cells. In their model the intracellular matrix is built from a set of stored expression patterns with the projection rule of Hopfield networks. This makes it symmetric, and the tissue settles into local minima of a global energy. We extend the model to directed gene regulation, for which no global energy exists.

Consider a tissue of *N* cells, each with the same *M* genes. The state of gene *A* in cell *i* is a binary spin,

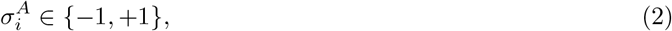

where +1 denotes an expressed gene and −1 a gene that is not expressed. The state of the full system can be read in two ways:

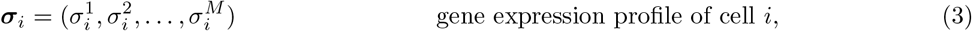

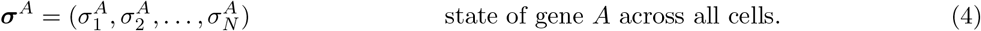

The model has two types of interaction.

#### Intracellular layer (gene regulation)

The *M* × *M* matrix *K* encodes the GRN, which is the same in every cell. *K*_*AB*_ is the effect of gene *B* on gene *A*: +1 for activation, −1 for repression, and 0 if there is no edge. Regulation is directed, so *K* is nonreciprocal,

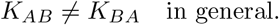

#### Intercellular layer (tissue structure)

The *N* × *N* adjacency matrix *J* records which cells communicate: *J*_*ij*_ = 1 if cells *i* and *j* are neighbours and *J*_*ij*_ = 0 otherwise, with *J*_*ii*_ = 0. Communication between cells is taken to be undirected, so *J* is symmetric,

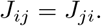

**Table 1.**
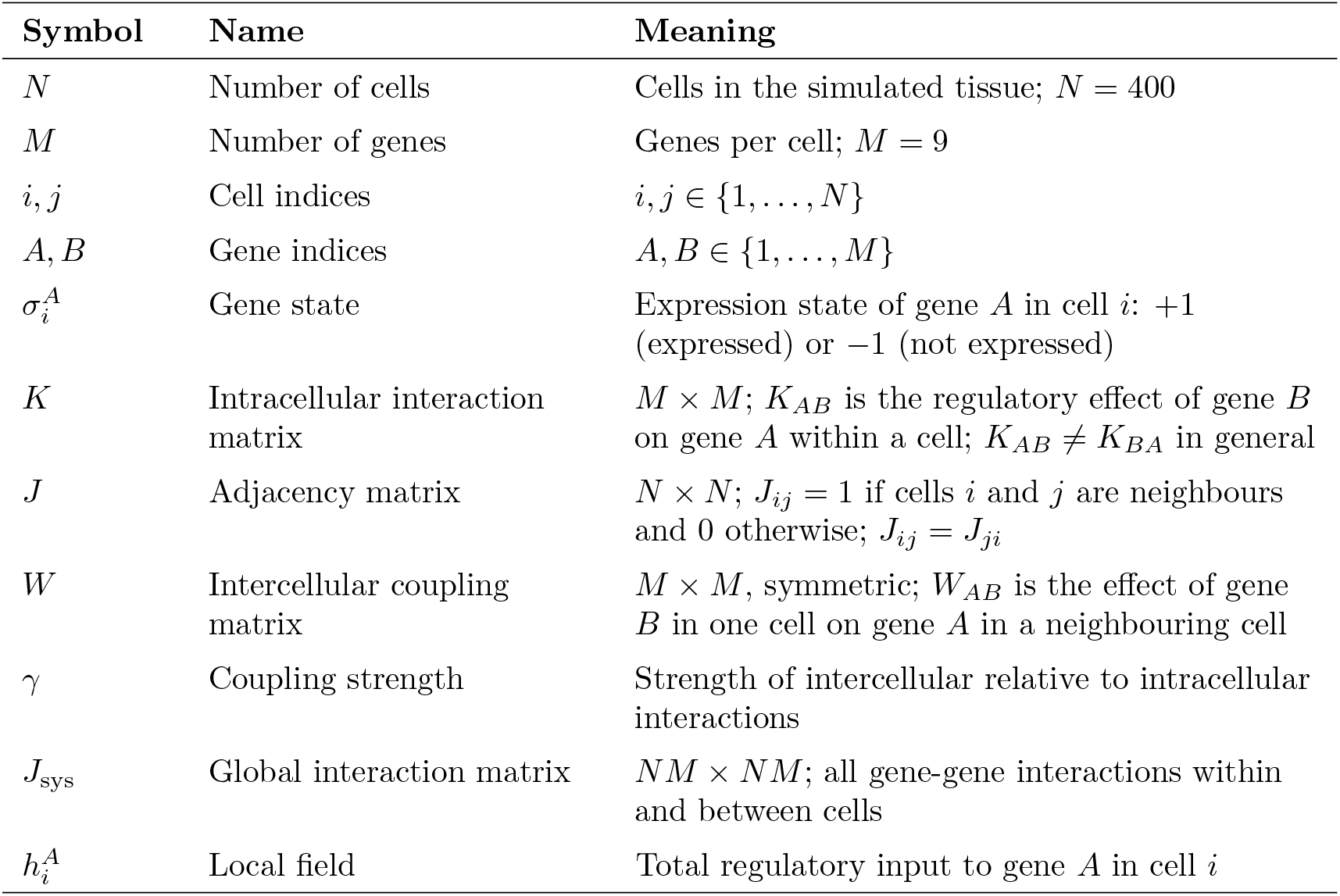
Variables of the nonreciprocal multicellular Ising model.

| Symbol | Name | Meaning |
| --- | --- | --- |
| $N$ | Number of cells | Cells in the simulated tissue; $N = 400$ |
| $M$ | Number of genes | Genes per cell; $M = 9$ |
| $i, j$ | Cell indices | $i, j \in \{1, \dots, N\}$ |
| $A, B$ | Gene indices | $A, B \in \{1, \dots, M\}$ |
| $\sigma_i^A$ | Gene state | Expression state of gene $A$ in cell $i$: +1 (expressed) or $-1$ (not expressed) |
| $K$ | Intracellular interaction matrix | $M \times M$; $K_{AB}$ is the regulatory effect of gene $B$ on gene $A$ within a cell; $K_{AB} \neq K_{BA}$ in general |
| $J$ | Adjacency matrix | $N \times N$; $J_{ij} = 1$ if cells $i$ and $j$ are neighbours and 0 otherwise; $J_{ij} = J_{ji}$ |
| $W$ | Intercellular coupling matrix | $M \times M$, symmetric; $W_{AB}$ is the effect of gene $B$ in one cell on gene $A$ in a neighbouring cell |
| $\gamma$ | Coupling strength | Strength of intercellular relative to intracellular interactions |
| $J_{\text{sys}}$ | Global interaction matrix | $NM \times NM$; all gene-gene interactions within and between cells |
| $h_i^A$ | Local field | Total regulatory input to gene $A$ in cell $i$ |

Cells sit on a 20 × 20 square lattice with periodic boundary conditions, so *N* = 400, and each cell has *M* = 9 genes. Each cell communicates with its eight nearest neighbours (Moore neighbourhood): the cells to its north, north-east, east, south-east, south, south-west, west, and north-west. These local interactions are a simple representation of short-range cell-cell communication, such as Notch or paracrine signalling.

Genes in neighbouring cells interact through the intercellular coupling matrix *W*. Its entry *W*_*AB*_ gives the sign and strength of the effect of gene *B* in one cell on gene *A* in a neighbouring cell. Following [49], *W* is symmetric, *W*_*AB*_ = *W*_*BA*_, with entries drawn independently from the uniform distribution on [−1, 1]. The same *W* applies to every pair of neighbouring cells and stays fixed during a simulation. Therefore, for the following results asymmetry only enters the model only through intracellular interactions *K*.

Combing intracellular and intercellular layers by summing Kronecker product gives the global interaction matrix:

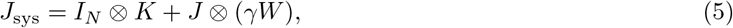

Acting on the tissue state vector **x** = vec(***σ***) with entries 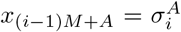. The vector **x** lists the state of every gene in every cell, so *J*_sys_ is an *NM* × *NM* matrix. The Kronecker product *X* ⊗ *Y* is a block matrix whose block (*i, j*) is *X*_*ij*_*Y*. Thus *I*_*N*_ ⊗ *K* places the GRN on the diagonal block of each cell, and *J* ⊗ *γW* places *γW* on the block of each pair of neighbouring cells [49]. The parameter *γ* ≥ 0 sets the strength of intercellular communication relative to intracellular regulation. Although *J* and *W* are symmetric *J*_sys_ is not, because *K* is not. Appendix **??** discusses the symmetric case, which corresponds to an undirected GRN.

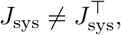

Without a global energy function, each gene follows an energy of its own, the **selfish energy** [49, 67]. The selfish energy of a gene depends only on its regulatory inputs, from within its cell and from its neighbours. A global Hamiltonian is not defined because there is no well-defined energy minima for non-equilibrium systems [23, 67] making the metric unreliable for describing the effects of GRN network dynamics on the formation of tissue structures. The total input to gene *A* in cell *i* is its local field,

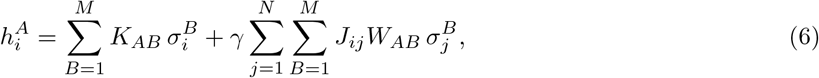

or **h** = *J*_sys_ **x** in vector form. The selfish energy of the gene is 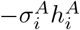, which is lowest when the gene agrees with the sign of its field. We write

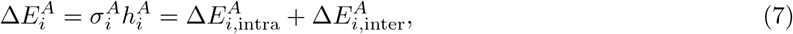

where

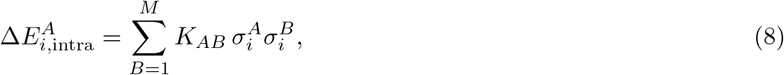

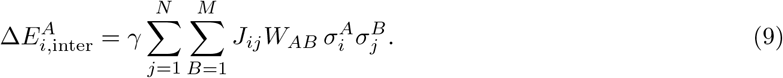

A flip of 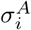 changes the selfish energy of gene *A* by 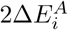, therefore lowering the energy when 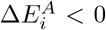 (when the gene state disagrees with the sign of its field). The intracellular term collects the inputs to gene *A* from the GRN of its own cell. The intercellular term collects its inputs from neighbouring cells, through *J* and *W*.

In terms of the global interaction matrix (Eq. (5)), the same quantity reads

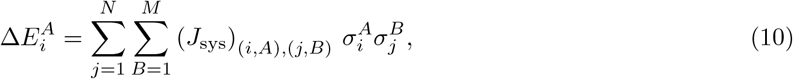

where the index pair (*i, A*) denotes row (*i* − 1)*M* + *A* of *J*_sys_.

#### Dynamics

Genes are updated one at a time with Glauber dynamics at zero temperature [72]. A gene flips if 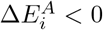 and keeps its state if 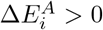. If 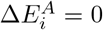, a gene in state (−1) flips to (+1) with probability 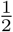, whereas a gene in state (+1) remains unchanged. Each update therefore aligns the gene with the sign of its local field, with ties broken at random. Because *J*_sys_ is not symmetric, the tissue need not reach a fixed point and can settle into sustained fluctuations. Appendix **??** gives the update scheme and the criteria for stopping a simulation.

#### Isolated cells

Setting *γ* = 0 removes all intercellular terms. Each cell then evolves under its own GRN alone, with 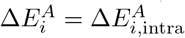 (Eq. (8)), and cells are independent. We refer to this as the isolated case.

### 2.4 Frustration metrics

Frustration measures how far a state is from satisfying its interactions. When the interactions of a network are mutually incompatible, no state satisfies all of them, and the network cannot settle into a state free of conflict. The pattern of unsatisfied interactions then carries information about the dynamics and stability of the system [47, 73]. The concept has been extended to biological networks, for example to study how they respond to perturbations [74, 75]. Genes within and between cells can make conflicting demands, for example through competition for resources or binding factors. We therefore define metrics that count the regulatory interactions of a cell that are not satisfied.

An interaction is satisfied when the states of its two genes follow its sign, and frustrated otherwise [47]. A satisfied interaction lowers the selfish energy, and a frustrated one raises it. For an edge of the GRN in cell *i*,

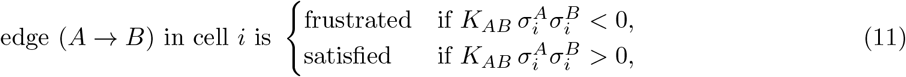

and there is no edge when *K*_*AB*_ = 0. Edges between cells are classified in the same way, by the sign of 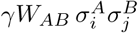.

We define two families of single-cell metrics (Figure 1). Intracellular frustration counts the frustrated edges of a cell’s own GRN, *K*. Intercellular frustration counts the frustrated edges between the genes of a cell and those of its neighbours, which are weighted by *γW*. Both are evaluated on the same tissue state, which the dynamics produce under the full matrix *J*_sys_. Together they split *J*_sys_ into its intracellular part, *I*_*N*_ ⊗ *K*, and its intercellular part, *J* ⊗ *γW*.

**Figure 1:**
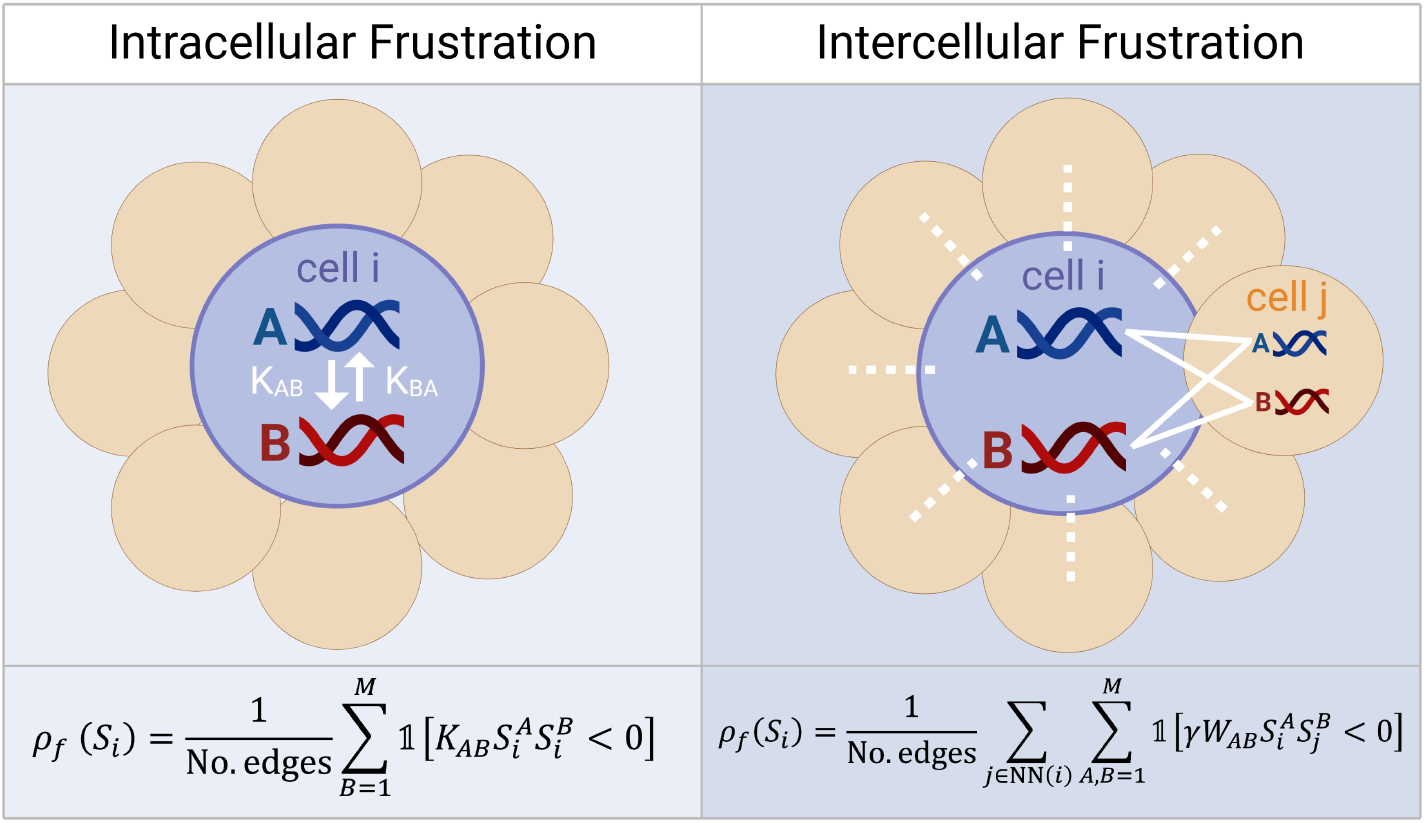
Intracellular and intercellular frustration. Cell *i* is shown in blue and its neighbours in beige; genes *A* and *B* are shown as DNA fragments. Left: intracellular frustration 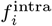 (Eq. (12)) counts the frustrated edges of the GRN within cell *i*. The arrows show the directed interactions *K*_*AB*_ and *K*_*BA*_, which differ in general. Right: intercellular frustration *f* ^inter^ (Eq. (15)) counts the frustrated edges between genes in cell *i* and genes in a neighbouring cell *j* (white lines), defined by the coupling *γW*. Edges within cell *i* do not contribute because *J*_*ii*_ = 0. Both metrics are evaluated on the same tissue state, which the dynamics produce under the full interaction matrix *J*_sys_.

#### Intracellular frustration

The intracellular frustration index of cell *i* is the number of frustrated edges in its GRN,

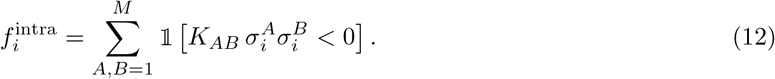

The intracellular frustration density divides this number by the number of edges in the GRN, *n*_*K*_, which is the number of non-zero entries of *K*:

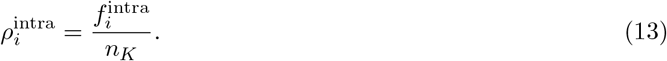

#### Isolated frustration

The isolated frustration density is the intracellular frustration density in the isolated case,

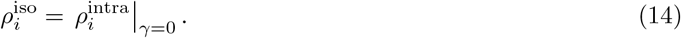

The two metrics use the same formula and differ in the states on which they are evaluated. For *γ >* 0 the state of a cell depends on its neighbours, so intracellular frustration carries a spatial imprint of the tissue. Isolated frustration reflects the GRN alone.

#### Intercellular frustration

The intercellular frustration index of cell *i* counts the frustrated edges between its genes and those of its neighbours NN(*i*),

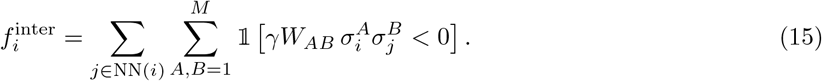

Because *J*_*ij*_ = 1 for *j* ∈ NN(*i*) and 0 otherwise, the sum over neighbours equals the sum over all cells weighted by *J*_*ij*_. Edges within cell *i* do not contribute because *J*_*ii*_ = 0. For *γ >* 0 the factor *γ* does not change the sign of any term, so *γ* affects this index only through the states. The intercellular frustration density divides the index by the number of intercellular edges of the cell,

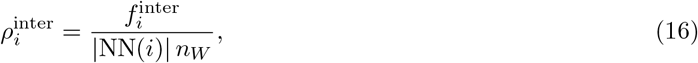

where *n*_*W*_ is the number of non-zero entries of *W*. For a dense *W* and eight neighbours the denominator is 8*M* ^2^ = 648. At *γ* = 0 there are no intercellular edges, and we set 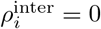.

### 2.5 Exemplar gene regulatory networks

We compare six directed GRNs, each with nine genes (Figure 2A). Genes are numbered top to bottom and left to right. The networks share the same downstream genes and differ only in the interactions among genes 1, 2, and 3.

**Figure 2:**
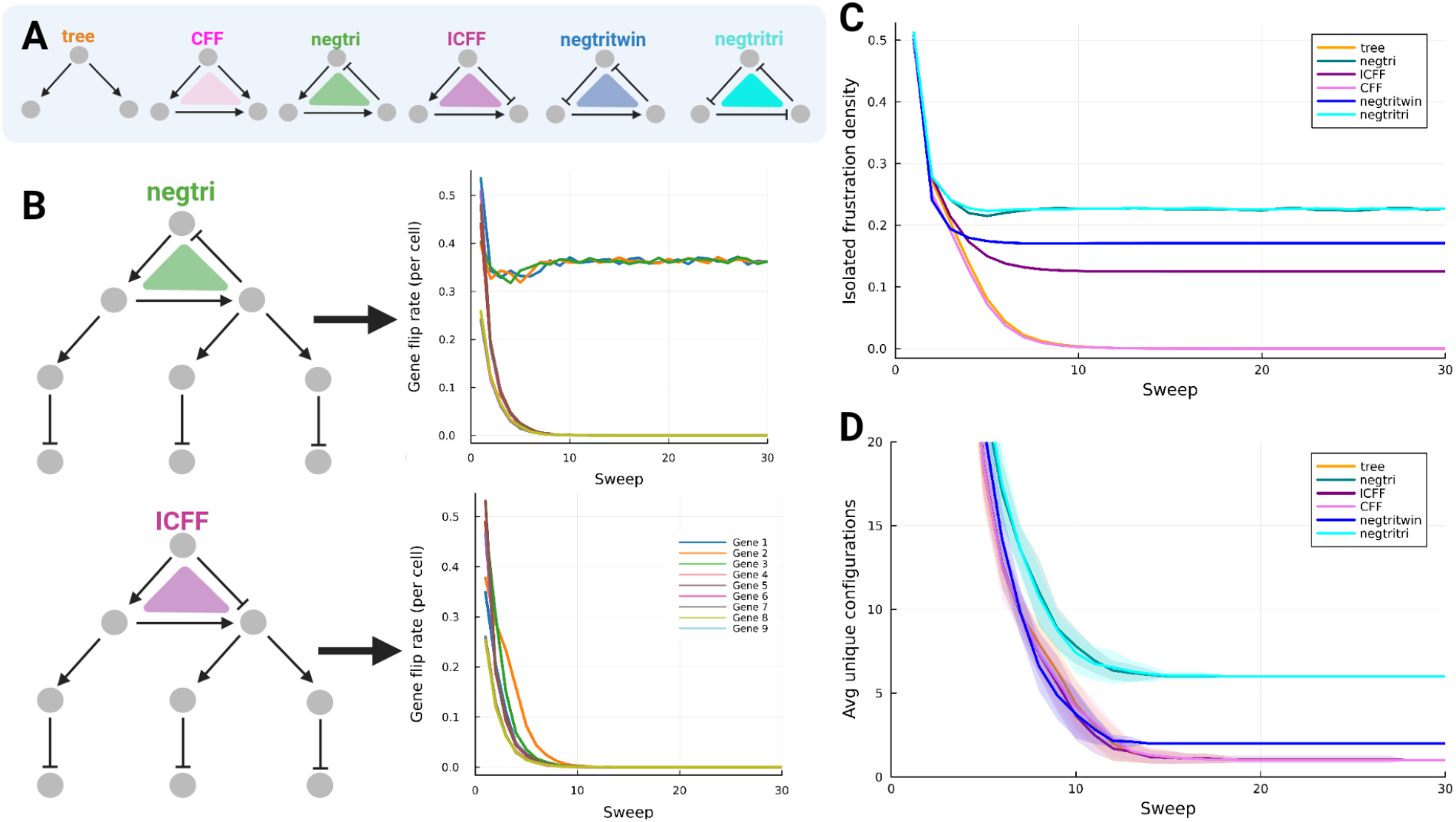
Frustration of isolated cells (*γ* = 0) for the six toy GRNs. (A) The loops among genes 1, 2, and 3 that distinguish the networks (Section 2.5). Arrows denote activation and bars repression. (B) The full nine-gene networks of negtri and ICFF, with the flip rate of each gene over the first 30 sweeps. (C) Isolated frustration density, averaged over the cells of the lattice, over the first 30 sweeps. (D) Number of distinct gene expression profiles across the lattice over the first 30 sweeps. Shading shows the range across simulations.

The loops among these three genes are of two types: In a *cyclic loop* each node has one incoming and one outgoing edge within the loop, so information flows around the loop in one direction. In a *non-cyclic loop* one node has two outgoing edges and another has two incoming edges; this part of the structure of coherent and incoherent feedforward loops.

The tree network has no loops: gene 1 activates genes 2 and 3. In CFF, gene 1 activates genes 2 and 3 and gene 2 activates gene 3, which forms a coherent feedforward loop. ICFF differs from CFF in that gene 1 represses gene 3, which makes the loop incoherent. The remaining three networks contain the cyclic loop 1 → 2 → 3 → 1, with one (negtri), two (negtritwin), or three (negtritri) repressive edges. By the sign rule for feedback loops, negtri and negtritri contain negative loops and negtritwin contains a positive loop [45].

All loops among genes 1, 2, and 3 have three nodes and three edges. Keeping the loop size fixed lets us compare motifs without the confounding effects of network size and connectivity, which become harder to separate once cells are coupled into a tissue.

### 2.6 Simulations

We simulate each network on the lattice of Section 2.3, starting from random initial states. For each network we draw 1000 realizations of *W* and run 50 simulations per realization at *γ* = 1 (Section 3.2) and 10 per realization when we vary *γ* (Section 3.3). At *γ* = 1 each intercellular coupling has the same scale as an intracellular one. Each simulation runs for at most 10,000 sweeps and stops earlier if the tissue freezes or settles into steady fluctuations. Appendix **??** gives the update scheme, the stopping criteria, and all settings (Table **??**).

## 3 Results

### 3.1 Isolated single cells

We first study how each GRN motif affects cells in isolation. Setting *γ* = 0 removes all communication between cells, so each of the 400 cells on the lattice evolves under its own GRN (Section 2.3). For each network we follow the isolated frustration density (Eq. (14)) and the number of distinct gene expression profiles across the lattice (Figure 2).

Negative cyclic loops produce the highest isolated frustration. Both negtri and negtritri settle at a density of about 0.23 (Figure 2C). In a negative loop at least one edge is frustrated in each state. When a gene in the loop flips to satisfy one of its edges, the frustration passes to the next edge. The frustrated edge therefore travels around the loop, and the genes of the loop keep flipping at a constant rate (Figure 2B, negtri). This argument holds for a single loop. Loops that share nodes can have different frustration properties [45, 47]. The network negtritwin, whose loop is positive, settles at a lower density of about 0.17.

Feedforward loops can also be frustrated. In ICFF, gene 3 receives activating signals from gene 2 and repression from gene 1. Once gene 2 follows gene 1, these inputs cancel, and one of the two edges into gene 3 stays frustrated whatever its state. Gene 3 has no edge back to genes 1 or 2, so the frustration does not propagate, and the network freezes (Figure 2B, ICFF). ICFF therefore settles at a non-zero density of about 0.13, below the cyclic loops. In tree and CFF every edge can be satisfied, and the density decays to zero.

The number of distinct profiles follows the frustration (Figure 2D). The lattices of negtri and negtritri hold about six profiles after 30 sweeps, and those of the other networks one or two. Because the genes of a negative loop keep flipping, isolated cells sit at different points of the same cycle, and the lattice holds several profiles at any time. Many profiles can share a frustration density, so the correspondence between the two measures is not exact.

Loop structure therefore sets how frustration arises and whether it persists. Cyclic loops pass frustration around the loop, and feedforward loops confine it to a single gene. These isolated values are the baseline for the coupled tissue.

### 3.2 Multicellular tissue

Cell-cell communication plays a central role in tissue formation and organization [49, 76]. Tissue structure emerges from cell–cell signalling acting on an underlying lineage tree [77]. Increasingly we can synthetically engieer such systems [78, 79]. Through cell-surface receptors and signalling molecules, such as Notch [50], cells exchange information that can change their intracellular regulatory dynamics. These dynamics shape transitions between cell states, often following complex stochastic differentiation trajectories [16, 61]. Inter-cellular signalling and intracellular regulation are therefore closely linked, and the collective behaviour of a tissue cannot be fully understood from isolated cells. We therefore couple the cells with *γ* = 1 and ask how communication changes the frustration of each cell. Figure 3 shows one final tissue for each GRN, its intracellular and intercellular frustration, and the distributions over all simulations.

**Figure 3:**
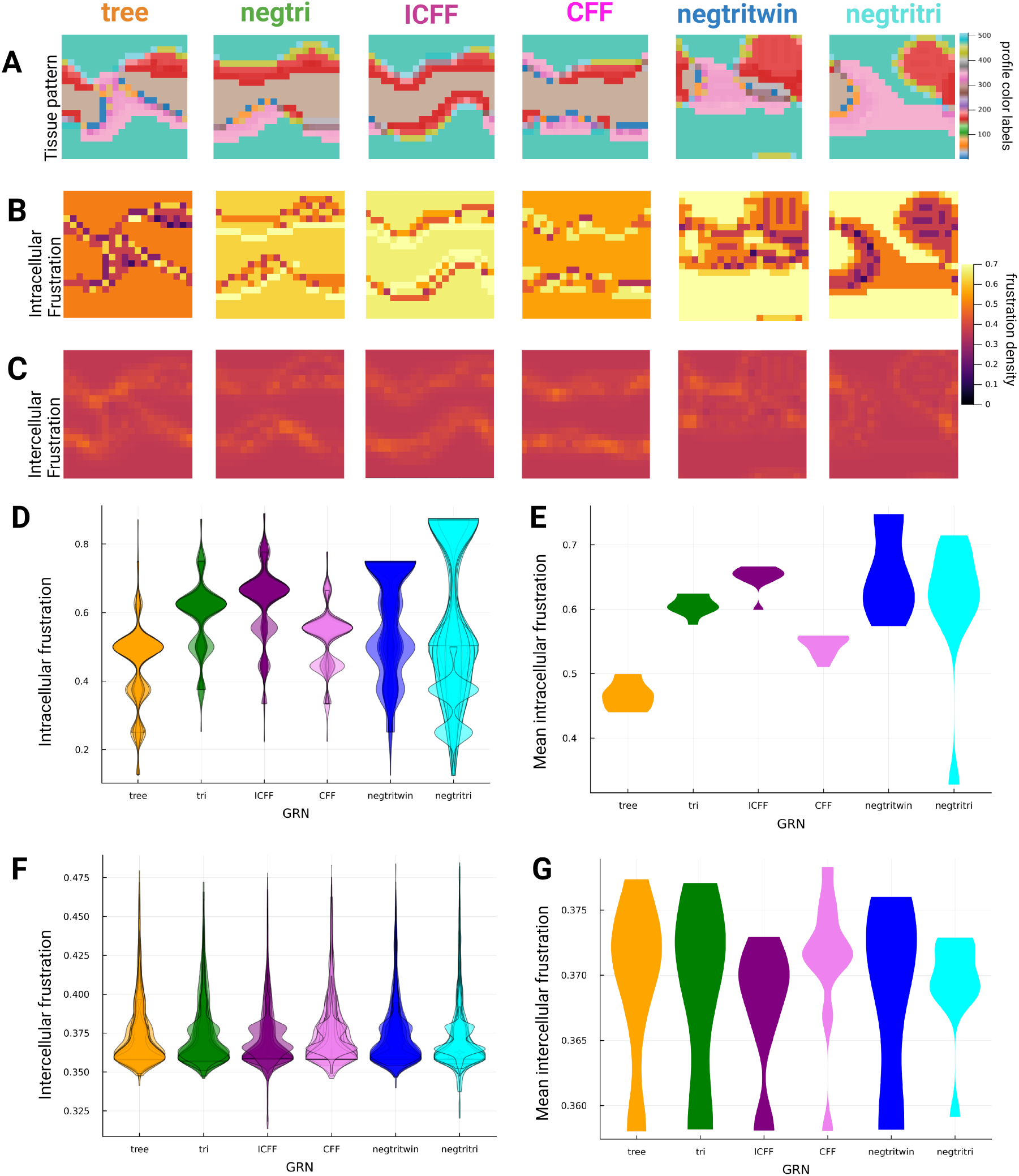
Frustration in a coupled tissue (*γ* = 1, realization *W*_2_). Panels A–C show the final tissue state from simulation 9. (A) Tissue expression pattern on the 20 *×* 20 lattice. Each colour corresponds to a distinct gene expression profile; the colour bar gives the profile index. (B) Intracellular frustration density of each cell (Eq. (13)). (C) Intercellular frustration density of each cell (Eq. (16)). (D) Intracellular frustration density distributions over all cells of the final tissue, with all 50 simulations overlaid as individual violins. (E) Mean intracellular frustration density per final tissue across the 50 simulations; each violin represents the distribution of per-tissue means. (F) As D, for intercellular frustration density. (G) As E, for intercellular frustration density.

Intracellular frustration depends on the gene expression profile of a cell (Eq. (12)). Cells of the same profile therefore share a value. Note more than one expression profile can result in the same frustration density (Figure 3A,B). Intercellular frustration depends on the profiles of neighbouring cells (Eq. (15)). Because intercellular interactions *W*_*AB*_ are symmetric, cells within a cluster exhibit low intercellular frustration densities. Cells at the boundary of a cluster experience competing intercellular interactions, resulting in an increase in intercellular frustration densities (Figure 3C).

Both frustration metrics are state-dependent: the gene expression profile of each cell *σ*_*i*_ evolves under the full global interaction matrix *J*_sys_, encoding the combinatorial effects of both intracellular and intercellular interactions. This is why intracellular frustration, which evaluates only 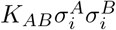 (Eq. (13)), still produces a spatial pattern that reflects the tissue structure (Figure 3A,B). Intracellular interactions *K*_*AB*_ are asymmetric — the directed regulatory influence of gene *B* on gene *A* is not reciprocated, meaning no single expression profile is guaranteed to satisfy all interactions simultaneously. One may interpret this as the spin state, updated with respect to *J*_sys_, satisfying or conflicting with *K*_*AB*_ depending on the directed structure of a GRN — producing a richer and more variable range of frustration densities at cluster boundaries compared to the intercellular metric.

For cells within a cluster, the magnitude of intracellular frustration corresponds to the GRN-inherent frustration established in the isolated single cell case study (Figure 2C): GRNs without inherent frustration — tree and CFF — produce lower intracellular frustration densities within clusters, while GRNs with inherent frustration — tri, ICFF, negtritwin and negtritri — produce higher densities, as seen in Figure 3D,E. Note that Figure 3 represents a single final tissue structure.

Coupling raises intracellular frustration for every GRN. The mean intracellular density is 0.46 for tree, 0.53 for CFF, and 0.60 to 0.65 for the networks with loops (Figure 3E), compared with 0 to 0.23 in isolation (Figure 2C). A random state gives a density of 0.5, because each edge is frustrated with probability 1*/*2. Each gene has potentially 8*M* = 72 intercellular inputs and at most three intracellular ones. Therefore at *γ* = 1 the state of a cell is set mostly by its neighbours.

The dynamics that we investigate here show that the interplay between intra-cellular and intercellular interactions is resolved globally at the tissue level. The ordering of the networks persists in the tissue. Tree and CFF, which have no frustration in isolation, have the lowest intracellular frustration (Figure 3D,E). The networks with loops have higher values, and tritwin and tritri vary most between simulations. Intercellular frustration shows no such separation, with means between 0.36 and 0.38 for every network (Figure 3F,G). The intercellular edges are the same for all networks and outnumber the intracellular ones, so intercellular frustration is set mainly by *W*. Note that a given frustration density value is a sum over many interactions (Eq. (13), (16)) — many combinations of satisfied and frustrated interactions can produce the same density value. Because each color in the tissue heatmap corresponds to a unique gene expression profile, tissues with identical frustration distributions need not correspond to the same spatial arrangement of expression profiles. Cluster size and number are therefore more biologically relevant quantities, as the spatial coordinates of such clusters are arbitrary under periodic boundary conditions.

### 3.3 Dissolving the tissue

We have coupled cells through cell-cell communication. As *γ* → 0 spatial cell-cell communication is turned down and ultimately off. We can use this to study the effect that cell-cell communication has on resolving frustration imposed by the GRN. Tracking intracellular and intercellular frustration as *γ* decreases shows how the conflict between a cell’s own GRN and its neighbours evolves as communication is dissolved.

As *γ* decreases, intracellular frustration decreases and (notional) intercellular frustration rises (Figure 4D,G). At high *γ* the state of each cell is set by its neighbours, and a cell’s frustration, imposed by its GRN, becomes less pronounced.

**Figure 4:**
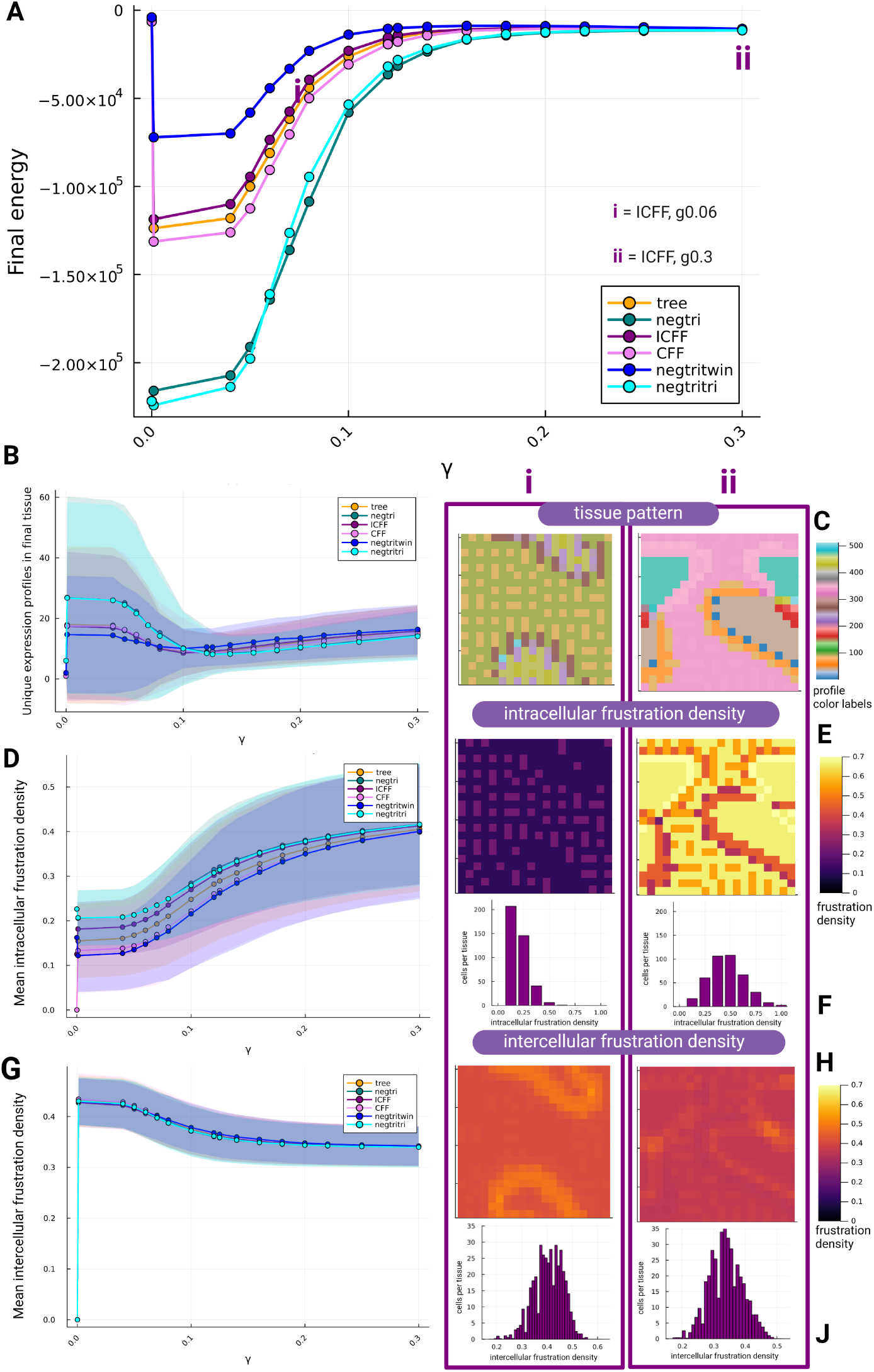
Dissolving the tissue by lowering *γ*. Results for 400 cells with 9 genes, 10 simulations per realization, and 1000 independent realizations of the random symmetric matrix *W*. Shading in B, D and G shows the standard deviation across realizations of *W*. (A) Final energy of the tissue, averaged over simulations and realizations of *W*. (B) Number of distinct expression profiles in the final tissue. (C) Final tissues for ICFF at (i) *γ* = 0.06 and (ii) *γ* = 0.3, from simulation 10 with realization *W*_2_. (D) Mean intracellular frustration density: the mean over the cells of each final tissue, averaged over all simulations and realizations of *W*. (E) Intracellular frustration density of each cell in the tissues of C. (F) Distribution of intracellular frustration density over cells, for all simulations with realization *W*_2_ at the values of *γ* in C. (G) Mean intercellular frustration density, defined as in D. (H, J) As E and F, for intercellular frustration density.

The change is most pronounced for *γ* ≈ 0.10. The final energy falls steeply below *γ* ≈ 0.12 (Figure 4A), and the number of distinct profiles is lowest near *γ* = 0.125 (Figure 4B). This is no surprise as for *γ* = 0.125, the eight neighbours, each coupled with weight *γ*, have together the same weight as a single intracellular edge. Below this value the intracellular term (Eq. (8)) dominates the selfish energy, and the intercellular term (Eq. (9)) becomes negligible *γ* → 0.

The two example ICFF tissues show the two regimes (Figure 4C,E,F,H,J). With weak coupling (i) the tissue becomes a mosaic of small patches; now intracellular frustration is concentrated below 0.25, and intercellular frustration is centered near 0.42. With strong coupling (ii) the tissue forms large clusters, intracellular frustration is spread around 0.45, and intercellular frustration is centered around 0.35. Below the transition (*γ* ≲ 0.125) each cell follows its own GRN with a weak bias from its neighbours. These two tissues are examples for a single GRN; across all six GRNs, the number of unique expression profiles per tissue in this regime separates into groups depending on the GRN, and the shading, which shows the standard deviation across the 1000 realisations of *W*, widens for every GRN (Figure 4B). The widening is most pronounced for negtri and negtritri, which in isolation (*γ* = 0) are unsatisfiable (Appendix 1). This describes the isolated GRN only: once cells are coupled, intra- and intercellular interactions combine in *J*_sys_, and a tissue may be satisfiable through combinations of these interactions even where the isolated GRN is not. The structure of a GRN constrains the expression profiles available to a cell, but because the dynamics are nonequilibrium and the resulting landscape is rugged, a cell can become trapped in a minimum and never explore all of them, so the profile it ends in depends on the realisation of *W* and on the initial condition. One may interpret the wider shading as this dependence becoming more visible as the asymmetric component becomes predominant. Since frustrated asymmetric GRNs may have no steady state, we describe these tissues as quasi-stable.

Below the transition the intracellular frustration of the networks separates (Figure 4D). At *γ* = 0 the values equal those of the isolated case (Figure 2C): negtri and negtritri are highest, followed by negtritwin and ICFF, with tree and CFF at zero. Intercellular frustration does not separate the networks at any *γ* (Figure 4G). As at *γ* = 1, the intercellular edges are the same for every network and outnumber the intracellular ones. At *γ* = 0 there are no intercellular edges, and intercellular frustration is zero by definition.

## 4 Conclusion

We have developed a nonreciprocal multicellular Ising model of gene regulation in tissues. The model keeps the direction of each regulatory interaction within a cell and couples neighbouring cells through random, symmetric signalling. Without a global energy function, each gene follows its own selfish energy, and the tissue need not settle into a fixed state. We defined intracellular and intercellular frustration densities to measure the conflicts that this creates within cells and between them.

In isolated cells the loop motif sets the frustration. Negative feedback loops cannot satisfy all of their edges, so the frustrated edge moves around the loop and the genes in the loop keep switching. The incoherent feedforward loop holds its frustration at a single gene, and the network freezes into a frustrated state. The tree and coherent feedforward networks reach states that satisfy every interaction. Frustration can therefore persist in two forms: it moves in a cyclic loop and stays in place in a feedforward loop. Populations of isolated cells with negative feedback loops hold the most distinct profiles, because their cells sit at different points of the same cycle.

Coupling cells into a tissue moves frustration between levels. Coupled cells form spatial clusters that share an expression profile, and the conflict between neighbouring cells concentrates at the boundaries between clusters. Stronger coupling lowers intercellular frustration and raises intracellular frustration, so cells satisfy more of the couplings to their neighbours and fewer of their own regulatory interactions. The tissue therefore resolves frustration only in part. In this sense cells compromise: they give up agreement with their own network to agree with their neighbours. The network still leaves an imprint, as networks with frustrated motifs remain more frustrated within the tissue than the tree and coherent feedforward networks.

## 5 Data & script availability

All original code has been deposited on the project GitHub repository https://github.com/taywood1123/Ising_on_top and is publicly available as of the date of publication. Julia 1.11.5 is used for all simulations, analyses, and visualizations. Any additional requests for data or code will be fulfilled by the lead contact upon reasonable request.

## Acknowledgments

The authors gratefully acknowledge the comments and suggestions from Augustinas Sukys, which have substantially improved the work and its presentation. We thank the members of the Theoretical Systems Biology group for their support and enthusiasm.

Both authors acknowledge the financial support through an ARC Laureate Fellowship to MPHS (FL220100005).

